# Combined short and long-read sequencing reveals a complex transcriptomic architecture of African swine fever virus

**DOI:** 10.1101/2020.07.18.202820

**Authors:** Gábor Torma, Dóra Tombácz, Zsolt Csabai, Norbert Moldován, István Mészáros, Zoltán Zádori, Zsolt Boldogkői

## Abstract

African swine fever virus (ASFV) is a large DNA virus belonging to the *Asfarviridae* family. Despite its agricultural importance, little is known about the fundamental molecular mechanisms of this pathogen. Understanding of genetic regulation provides new insights into the virus pathogenicity, which can help prevent epidemics. Short-read sequencing (SRS) is able to produce a huge amount of high-precision sequencing reads for transcriptomic profiling, but it is inefficient for the comprehensive annotation of transcriptomes. Long-read sequencing (LRS) is able to overcome some of the limitations of SRS, but they also have drawbacks, such as low-coverage and high error rate. The limitations of the two approaches can be surmounted by the combined use of these techniques. In this study, we used Illumina SRS and Oxford Nanopore Technologies LRS platforms with multiple library preparation methods (amplified and direct cDNA sequencings and native RNA sequencing) for constructing the transcriptomic atlas of ASFV. This work identified a large number of novel genes, transcripts and RNA isoforms, and annotated the precise termini of previously described RNA molecules. In contrast to the current view that the ASFV transcripts are monocistronic, we detected a significant extent of polycistronism. A multifaceted meshwork of transcriptional overlaps is also discovered.

## Introduction

High-throughput massively parallel sequencing has revolutionized modern biology by facilitating the explosive growth of genomics and transcriptomics. Next-generation short-read sequencing (SRS) is able to produce a tremendous volume of data with unprecedented speed. Currently, Illumina rules the SRS market. RNA studies are hampered by the limitations of SRS that cannot sufficiently cope with the complexity of large transcriptomes. Third-generation long-read sequencing (LRS) can overcome many of the drawbacks of SRS, which include the inefficiency to make a distinction between transcription isoforms, the low efficiency of detecting long RNA molecules, and transcriptional overlaps. The LRS platforms have two main disadvantages: the relatively low throughput and the high error rate. However, with the combined use of these technologies high quality and high throughput data can be produced on full-length transcripts.

African swine fever (ASF) is a highly lethal haemorrhagic disease of domestic pig. Its causative agent is the ASF virus (ASFV) that is the sole member of the *Asfarviridae* family [1]. The natural hosts of the virus are warthogs, bush pigs and soft ticks of the genus *Ornithodoros*, that makes ASFV the only DNA virus that can infect both mammals and arthropods [2]. To date, 24 genotypes of the virus have been identified [3]. ASFV transmitted from the natural hosts to swine in multiple waves. The first transmission most probably happened in Africa where the disease was first reported in 1921 [4]. Genotype I of ASFV spread to the South-European and Caribbean regions in the ‘60s and ‘70s, respectively [5], whereas genotype II emerged in Georgia in 2007, where it spread rapidly westward in Eastern Europe [6], later eastward, and now reached the major Asian swine producer countries, China and Vietnam [7, 8]. Since effective vaccine or antiviral drugs are unavailable against the virus at this moment, it unquestionably presents the largest world-wide economic threat to animal industry in recorded history.

ASFV belongs to nucleocytoplasmic large DNA viruses. Even though some form of nuclear replication is detected in the early phase of the infection [9], it mainly replicates itself in the cytoplasm. The genetic organization of ASFV best resembles to that of the poxviruses [10, 11]. The viral genome is ~170-190 kbp long and contains covalently closed terminal repeats in its ends. The variation between the genomes of viral strains mainly originates from the varying number of genes belonging to the multigene families (MGFs), that are located at the genome termini [10]. Genetic analyses have revealed over 160 ORFs in the viral genome, although the exact number of protein coding genes are not known yet. Proteomic analysis of highly purified extracellular ASFV particles identified 68 viral proteins [12]. Twenty ASFV genes are considered to be involved in the transcription and modification of the viral mRNAs [11]. The proteins of another 17 genes (~17 % of the genome) are involved in nucleotide metabolism, DNA replication, as well as DNA repair and modification [10, 13]. The members of the five MGFs (MGF 100, MGF 110, MGF 300, MGF 360, MGF 530) constitute another large group of ORFs (30-49 depending on isolates), and they occupy ~18-30% of the genome. MGF proteins are involved in immune evasions and are important virulence and host range factors [10, 13, 14]. The temporal regulation of ASFV gene expression appears to be similar to the transcription cascade described in poxviruses [11, 15], in which four classes of the consecutively transcribed RNAs have been described. The ASFV immediate early (IE) and early (E) genes are mainly synthesized before the DNA replication, whereas intermediate (I) and late (L) genes are transcribed after the onset of DNA replication [11].

The ASFV ORFs have been reported to be transcribed as monocistronic RNA molecules [11, 16, 17], which if so, represents a different principle of transcriptome organization than those of other DNA viruses, which express a large number of multigenic transcripts (e.g., poxviruses, herpesviruses, and baculoviruses; [18–23]. ASFV ORFs are compactly laid in both strands with very short intergenic regions or with minimal overlaps with tail-to-head, head-to-head or tail-to-tail arrangements [10, 13]. Regulatory elements and promoters usually overlap with upstream coding regions, hence their accessibility for the protein elements of the transcription complex, and consequently, their activity must be largely influenced by the transcription activity of the upstream gene. Very little information is available on the functioning of ASFV promoters. At late phase of infection, p30 and DNA polymerase promoter have higher activity than that of the p72 promoter regulating the transcription of the most abundant viral protein p72 [24]. The four-nucleotide core TATA sequence appears to be an essential motif in the majority of late promoters [25].

The transcription kinetics of ASFV has only been studied in a few cases in details. Transcripts are initiated and terminated at precise sites upstream and downstream of the ORFs, respectively [26]. The transcribed RNAs have 5’-cap structures and 3’-poly(A) tails with an average length of 33 nucleotides added by the viral capping enzyme complex and the poly(A) polymerase, respectively [27]. Jaing and coworkers have carried out a survey on the entire ASFV transcriptome using RNA-seq [28]. Approximately 60% of the ORFs (109 genes) were detected from the circulating monocytes of highly pathogenic Georgia 2007/1 infected pigs. Although this study has provided valuable information about highly expressed ASFV genes, it has not supplied sufficient data to construct a detailed transcriptional map of the virus. In a very recent study, Cackett and co-workers reported an SRS analysis on ASFV transcriptome [29]. The authors also carried out cap analysis gene expression sequencing (CAGE) and 3’-RNA-Seq.

Many wild-type ASFV strains cannot be propagated in established cell lines [30]. However, the virus replicates relatively well in porcine primary alveolar macrophages (PAMs) *in vitro*, although the sensitivity of naïve PAM culture to ASFV infection varies batch by batch [31]. In this study, we applied a dual sequencing approach for the investigation of ASFV transcriptome.

## Results

### Analysis of the ASFP transcriptome with a dual sequencing approach

In this work, we report the application of the combined use of SRS (Illumina MiSeq) and LRS (ONT MinION) techniques for the analysis of the ASFV transcriptome. We used anchored oligo(dT) primers in the LRS and random oligonucleotides in the SRS approach for priming the reverse transcription (RT) of the first cDNA strand. Both primers were linked to short adapter sequences in those samples which were amplified by PCR after the RT. We applied three ONT sequencing methods: cDNA sequencing with or without PCR amplification [the latter is termed direct cDNA sequencing (dcDNA-Seq)], and amplification-free native (direct) RNA (dRNA) sequencing. The Illumina SRS yielded altogether 69,068 reads mapped to the ASFV genome with an average genome coverage of 50.13. Illumina sequencing was primarily used for the validation of novel transcripts and for increasing the sequencing precision. The amplified cDNA-Seq produced 126,763 ASFV reads, with an average read length of 458 bps and genome coverage of 304,5. Direct cDNA sequencing generated 8,587 viral reads with an average read length of 568 bp. Native RNA sequencing yielded 4,361 viral read, with an average read length of 637 bp and with an average genome coverage of 14,6. We used the LoRTIA software suit, developed in our laboratory, for the identification of TSS and TES positions of the putative transcripts. The RT and the PCR may generate technical artefacts, such as nonspecific binding of oligod(T) or PCR primers and template switching. In addition to poly(A) tails, oligo(dT) primers may also hybridize to A-rich regions of RNA molecules and thereby produce spurious transcripts. Additionally, it has been shown that cDNA library preparation of ONT protocol induces read truncation for transcripts containing internal runs of T’s [32]. These false products were removed from further analysis with the help of LoRTIA tool, albeit in many cases we were unsure about the non-specificity of the discarded reads. We used strict criteria for further filtering the potential spurious transcript: only those reads were accepted as transcripts, which were detected in at least two independent experiments with precise transcript termini. We applied even stricter criteria for accepting ncRNAs and truncated RNAs: besides two independent samples, either dcDNA and/or dRNA technique also had to confirm these transcripts. In certain genomic regions, very low coverage was obtained, therefore the strict criteria were not applied for the annotation of TSSs and TESs (Fig. 1).

**Fig. 1.**
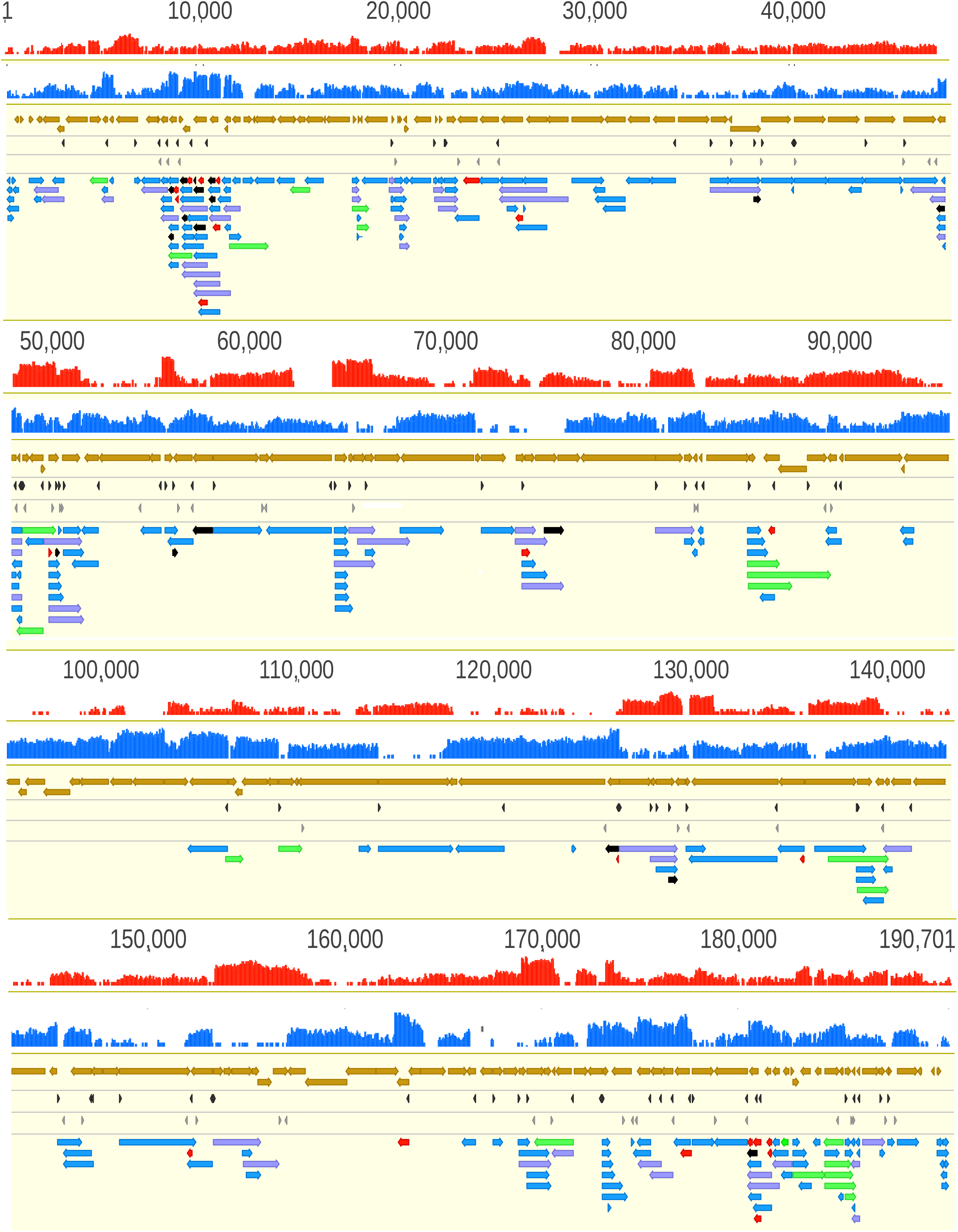
African Swine Fever Virus transcriptome. Location of the annotated ASFV transcripts on the viral genome. The red and the blue histograms represent the coverage of cDNA reads on the two DNA strands. The mRNAs have a higher coverage than the antisense RNAs, which, in principle, can be produced from their own promoters, or are a readthrough products of convergent genes, or are resulted by the TSS overlaps of divergently-oriented gene products. Colour code: light brown arrow-rectangles: coding sequences; dark grey arrow-rectangles: previously annotated TSS; light grey arrow-rectangles: previously annotated TES; black arrow-rectangles: 5’-truncated in-frame ORFs containing transcripts; purple arrow-rectangles: novel polycistronic transcripts; red arrow-rectangles: novel non-coding transcripts; while green arrow-rectangles: novel complex transcripts.

Altogether, we detected a total of 132 LoRTIA and 70 non-LoRTIA TSSs, as well as 137 LoRTIA and 83 non-LoRTIA TESs (Supplementary Table S1). In sum, we detected and annotated altogether 311 novel ASFV RNA molecules (Fig. 1, Supplementary Table S2), including 273 full-length transcripts with precisely identified termini and 38 RNA molecules without accurately annotated TSSs. We obtained 69.01 bp length for an average 5’-untranslated region (UTR) length, and 369.31 bp length for the average 3’-UTR lengths. We obtained the TSS of 92 and the TES of 57 earlier annotated TSSs and TESs [29].

### Comparison of the genome of strains ASFV_HU_2018 and Ba71V of ASFV

The genome size of the various ASFV strains can significantly vary [33]. We used ASFV_HU_2018 isolate ([31]; which is probably the same as strain Georgia 2007/1 [6]) and compared it to the strain Ba71V used for transcriptome analyis by Cackett and colleaguses in a very recent publication [29]. We obtained that Ba71V genome lacks 15,143 bps (22 genes), which is present in our ASFV_HU_2018 strain (Supplementary Table S1).

### Novel putative protein-coding genes

*5’-truncated transcripts with in-frame ORFs* It has recently been shown in several viruses that the transcripts containing 5’-truncated in-frame ORFs within larger canonical ORFs - of which they share stop codons - are much more common than it has previously been thought [21, 22, 34]. Here we demonstrate the same phenomenon in ASFV transcripts (Fig.2a). We compared our results with the CAGE analysis published by Cackett and colleagues [29] (Supplementary Table S3). In total, we discovered 19 novel putative mRNAs with shorter in-frame ORFs (Supplementary Table S2), of which 3 (A137R, CP312R, MGF 100-1L) have also been detected by Cackett and co-workers. Some of these TSSs were also present in Cackett and colleagues’ non-primary TSS list. Further experiments are needed to demonstrate whether this coding potential is realized in translation in every cases.

**Fig. 2.**
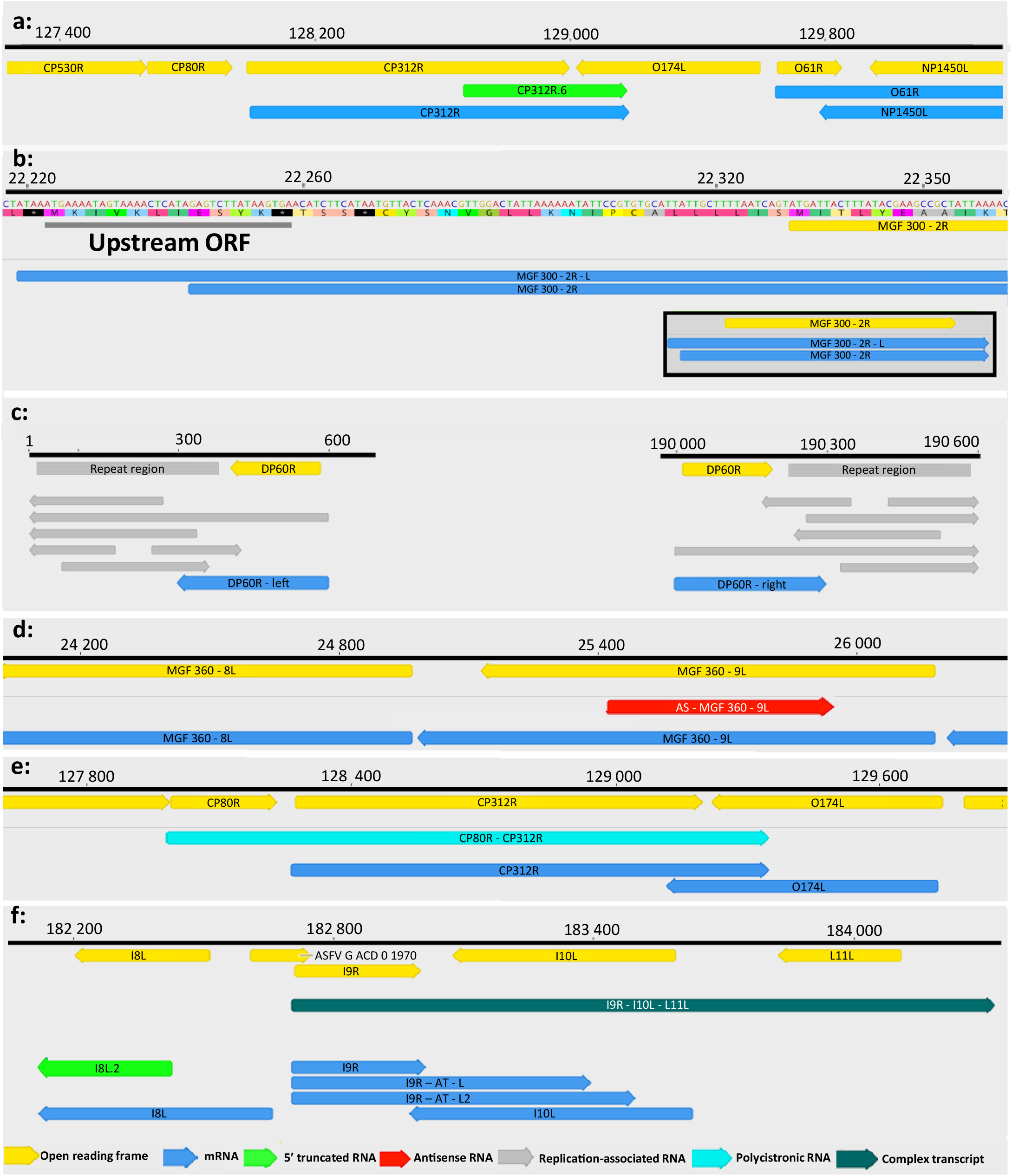
Examples for the classes of ASFV transcripts. (a) ***5’-truncated mRNAs*** This picture illustrates the mRNA type containing short 5’-truncated in-frame ORFs (CP312R.6; green arrow) within the canonical ORFs (CP312R; yellow arrow). (b) ***upstream ORF-containing transcripts*** MGF 300-2R transcript and its longer 5’-UTR isoform (MGF 300-2R-L) is illustrated in this picture. The grey line indicates the uORF sequence. The bracketed picture contains two transcript isoforms (blue arrows) and the ORF of the gene encoding them (yellow arrow). (c) ***replication-associated RNAs*** This figure illustrates the repeat region at genomic termini and the raRNAs overlapping the repeats (grey arrows). (d) ***antisense RNAs*** The red arrow represents an asRNA (AS-MGF 360-9L) overlapping the MGF 360-9L gene in opposite polarity. (e) ***polycistronic RNAs*** CP80R-CP312R bicistronic transcript (light green arrow) comprising two tandemly oriented genes. (f) ***complex transcripts*** The I9R-I10L-L11L complex transcript (dark green) comprises three genes of which I9R stands in an opposite orientation compared to the other two genes (L11L.2 and I10L).

*Novel intergenic transcripts with small ORFs* In this study, we detected four LoRTIA and three non-LoRTIA novel small RNA molecules containing short (9-192 bp) ORFs encoded in various intergenic regions (Supplementary Table S2). Cackett and colleagues have also identified one of these transcripts (pNG6). Further investigations have to be carried out to ascertain whether these small transcripts are translated.

### Upstream ORF-containing mRNAs

Recent advances in gene expression studies have shown that more than half of the RNA molecules contains translationally active uORFs located upstream positition relative to the canonical ORFs [35]. This study revealed in total 30 uORFs in the mRNAs, 6 of them overlapped with the main ORFs (Supplementary Table S2, Fig. 2b). Additionally, six of these uORFs were only present in the long TSS isoforms of transcripts but not in the short one. The average length of these ORFs are 42,7 bps. The average distance between the stop codon of uORFs and the ATG of canonical ORFS is 45,76 bps (Supplementary Table S1).

### Putative non-coding transcripts

Non-coding RNAs (ncRNAs) are specified by RNA genes that are located either at intergenic regions, or within protein-coding genes, and they can be encoded by both the positive and negative DNA strands of protein-coding genes. In this work, we detected several long ncRNAs (lncRNAs >200 bp in length), including inter- and intragenic transcripts, antisense RNA (asRNAs), and replication-associated RNAs (raRNAs).

*Intergenic transcripts* We detected three putative ncRNAs with an average length of 118 bp, which were located at intergenic genomic regions (Supplementary Table S2).

*The 3’-truncated transcripts* are controlled by the same promoters as the overlapping mRNAs but these embedded RNA molecules are terminated at sequences preceding the stop codons, therefore they do not contain ORFs. In ASFV, we detected 22 such transcripts. The LoRTIA tool was used for screening false priming on the A-rich regions of transcripts. Further studies have to be determined whether this group of transcripts is functional or only spurious RNA molecules.

*Antisense RNAs* can in principle be controlled by distinct promoters, or they can be generated by transcriptional readthrough from adjacent or distal convergent genes. In our LRS approach, we detected seven asRNA fragments with identified TESs, but unannotated TSSs (Fig. 2c). Our SRS approach however detected a pervasive asRNA expression along the entire ASFV genome. (Fig. 1). It is assumed that the majority of these transcripts are generated by occasional transcriptional read-throughs from the neighbouring convergent genes or by divergent overlaps of the 5’-UTR transcripts. We could not confirm the existence of any asRNA, which are controlled by distinct cis-regulatory sequences, however, we cannot exclude the existence of such transcripts.

*Replication-associated RNAs* overlap the replication origins. RaRNAs have been identified in every examined virus [36]. In herpesviruses, these transcripts can be non-coding or alternatively, the longer TSS or TES variants of mRNAs [37]. In ASFV raRNAs we detected 6 low-abundance raRNA molecules at late infection times (12 to 20 hours), which are encoded at the repeat region (Fig. 2d).

### Transcript isoforms

*TSS isoforms* contain the same ORFs, but differ in the length of their 5’-UTRs and are usually controlled by distinct promoters. In this study, we obtained a comparatively low diversity of TSSs for which the reason may be the relatively low data coverage and the strict criteria used for the identification of novel transcripts. We detected 16 TSS isoforms of which 14 were longer and 2 shorter than the canonical transcripts. Only a single TSS isoform (MGF 300-2R) was identified by Cackett and colleagues [29] (they detected altogether 8 TSS variants), but two of our isoforms were present in their CAGE-seq non-primary TSS list (D345L, C147L).

*TES isoforms* contain the same TSS, but have distinct poly(A) signals and 3’-end transcript terminals. In this work, we identified 220 TESs, and 49 TES isoforms. Some of these TESs (ASFV G ACD 00600-A224L-AT-L, A151R-AT-S2, I73R-AT-L4, I215L-AT-S,MGF 360-18R-DP71L-DP96R-AT-L) were also found in Cacket at el. [29] non-primary p(A)-Seq data.

This study did not identify any *spliced transcripts* of ASFV.

### Polycistronic transcripts

Until now, it has been widely assumed that ASFV genome encodes exclusively monocistronic transcripts [11, 16, 17]. Here we show that it is not the case, since our study revealed extensive polycistronism in the ASFV transcriptome. Altogether, 41 bicistronic, 8 tricistronic, and 2 tetracistronic transcripts have been detected using ONT sequencing techniques (Fig. 2e). Fourteen of these transcripts share their TESs with the first genes of the polycistronic unit that are also expressed as monocistronic RNA molecules. Our investigations revealed that the organization of ASFV transcriptome is more similar to those of the poxviruses than it was expected [38]. Additionally, we demonstrated that, due to the large number of coterminal transcripts, ASFV transcriptome also resembles to a certain extent to those of herpesviruses [23] and baculoviruses [21]. The herpesvirus tandem genes tend to form co-terminal poly(A) sites with the following pattern on e.g. a tricistronic transcripts, where ‘a’ is the most upstream and ‘c’ is the most downstream gene: abc, bc, and c. ASFV can also form abc tricistronic RNA molecules, but each gene can also be expressed individually. The longest tricistronic transcripts (MGF 360-11L-MGF 360-10L-MGF 360-9L) is 3,499 nt long.

### Complex transcripts

Our LRS study demonstrated that ASFV expresses complex transcripts, which, *per definitionem*, contain at least one gene with opposed polarity in a polygenic transcript. We identified in total 22 complex transcripts. (Fig. 2f).

### Transcriptional overlaps

Transcripts can form parallel, convergent and divergent overlaps. Polycistronic ASFV transcripts represent parallel overlap of RNA molecules encoded by tandem genes. The distinctive feature of ASFV convergent overlaps compared to herpesviruses is the large number of ‘hard’ overlaps (all transcripts overlap with the convergent transcripts for a gene pair) and the large fraction of overlaps in the ‘soft’ overlaps (only certain ratio of transcript forms overlaps as a result of transcriptional read-through). This genomic organization suggests an important role of the transcriptional interference in the regulation of genome-wide gene expression of ASFV [39]. We detected 540 parallel, 60 convergent and 19 divergent overlaps. The average lengths of overlaps are as follows, head-to-tail overlaps: 595.3 bp; tail-to-tail overlaps: 319.4 bp; and the head-to-head overlaps: ~325.1. The uncertainty of the size of divergent overlaps comes from the fact that most of them were detected by dRNA-Seq, which produces sequencing reads inherently missing 15-35 bp from the 5’-UTRs of the transcripts (Supplementary Table S1).

## Discussion

African swine fever virus is one of the most important animal pathogen causing severe losses in animal husbandry. Despite its economic significance, no effective therapy against this virus is available yet. Understanding the architecture of viral transcriptome is not solely an academic issue because structure-based drug development is essentially dependent on our knowledge about the fundamental molecular mechanisms.

Here we report the first long-read sequencing study on the ASFV transcriptome, which also applies a short-read sequencing approach. The combination of LRS and SRS techniques results in practically error-free, high-coverage, full-length transcription reads. This analysis revealed numerous novel genes, transcripts and transcript isoforms, and ascertained the transcription start and end positions for most of the ASFV transcripts. Additionally, in contrast to the earlier views that the ASFV expresses exclusively monocistronic transcripts [11, 16, 17], our work demonstrates the widespread expression of multigenic RNA molecules, including polycistronic and complex transcripts, along the entire viral genome.

Most of the novel putative genes are embedded into larger canonical genes and contain in-frame ORFs. The SRS approach is inefficient in the discovery of these genes; that is why they had gone undetected before. Another class of putative protein-coding genes that was detected in intergenic locations encodes relatively short transcripts with small ORFs. Transcripts containing upstream ORFs beside the canonical ORFs represent another class of RNA molecules. We identified a number of uORFs, and found that none of them are expressed on separate RNAs independently of the canonical ORFs. Although the importance of uORFs is currently not very well understood, it has been suggested that they may have an important influence on gene-expression [40–42]. Additionally, we demonstrated a potentially intriguing variance in a few cases; namely, that only the longer TSS isoforms of transcripts contain the uORF, but not the shorter ones. We observed a similar phenomenon in human cytomegalovirus [22]. Potentially, the two transcript isoforms might provide a distinct translational regulation of the same gene at different stages of viral infection. The protein-coding capacity for all type of novel ORFs has to be experimentally verified.

We also detected various classes of non-coding transcripts, including intra- and intergenic transcripts and antisense RNAs. The asRNAs are either transcriptional readthrough products of convergent genes, or are generated by divergent overlaps. No asRNAs with own promoters have been detected in this work. This study also identified several novel TSSs and TESs and confirmed many of the transcript termini published recently by others [29]. Additionally, we could not detect any spliced ASFV transcripts.

Polycistronic transcription is widespread in prokaryotic organisms and in certain viruses, but is rare in eukaryotes. In bacteria and bacteriophages, the Shine-Dalgarno sequences control the translation of downstream genes on multigenic transcripts, whereas some RNA viruses develeloped a variety of mechanisms to tackle this problem, which includes the use of internal ribosome entry site sequences, ribosomal frameshifting, or leaky ribosomal scanning. However, the function of polycistronism in large DNA viruses is not clearly understood, because the translation initiation in this organisms is cap-dependent, therefore, the downstream genes on multigenic transcripts are untranslated. Only a few exceptions to this rule have been discovered so far, which include the translation of uORFs in addition to the downstream canonical ORFs [43–44].

## Methods

### Cells, viruses and infection

Pulmonary macrophage (PAM) cells were harvested freshly from swine lungs according to the OIE Manual’s instructions [45]. PAM cells were used for the propagation of the highly virulent ASFV_HU_2018 isolate of African swine fever virus (ID Number: MN715134). PAM cells were grown in RPMI 1640 containing L-glutamine (Lonza) medium supplemented with 10% foetal bovine serum (Euro Clone), 1% Na-pyruvate (Lonza), 1% non-essential amino acid solution (Lonza), and 1% antibiotic-antimycotic solution (Thermo Fisher Scientific) at 37°C in 5% CO2 in air gas phase. The infectious titre of serially diluted viral stock was calculated using an immunofluorescence (IF) assay as described earlier [31]. PAMs were cultivated in 6-well plates at a density of 3.3×10^5^ cells and infected at a multiplicity of infection (MOI) of 10 plaque-forming unit per cell at 4 h after cell seeding. Supernatant was replaced with fresh medium after 1 h post-infection (p.i.). Infected PAM cells were harvested at 4, 8, 12, and 20 h p.i., whereas cock-infected cells were harvested at 20 h p.i. IF assay was used for monitoring the efficiency of infection in an infected control well fixed at 20 h p.i. Infection efficiency remained at approximately 20% despite the high viral titre (MOI = 10) applied for the infection. This result is in agreement with other’s observations.

### RNA purification

#### Extraction of total RNA

We used the NucleoSpin^®^ RNA (Macherey-Nagel) kit for isolation of the total RNA from the samples as was described in our previous publications [46]. Briefly, samples were incubated with a lysis buffer (supplied by the kit), then DNase I treatment was carried out. Purified RNA samples were eluted from the silica membrane in nuclease-free water. Samples were stored at −80°C until further use. The total RNAs were used directly for the “amplified cDNA protocol” from ONT.

#### Purification of polyadenylated RNAs

For the dRNA and dcDNA sequencing approaches the poly(A)+ fraction of the total RNAs were extracted. This process was carried out with the Qiagen’s Oligotex mRNA Mini Kit using spin columns according to the Kit’s manual.

#### Removal of the ribosomal RNAs

The RiboMinus™ Eukaryote System v2 (Thermo Fisher Scientific) was used to obtain rRNA-free RNA samples which is required by the applied Illumina library preparation approach. The ribodepletion was carried out according to the kit’s instructions.

### Library preparations

#### Direct RNA sequencing on MinION SpotON Flow Cells

Native RNA sequencing was carried out using the ONT SQK-RNA002 kit as was described earlier [38, 47]. In brief, PolyA(+) RNA mixture (including 4, 8, 12, and 20 h p.i.) was reverse transcribed using oligo(dT) primer (from the ONT kit), NEBNext Quick Ligation Reaction Buffer, and T4 DNA ligase, dNTPs (both from New England Biolabs), and enzyme and buffer from the Invitrogen’s SuperScript III kit. RNA-cDNA hybrids were washed with Agencourt RNAClean XP Beads (Beckman Coulter). ONT RMX adapters were ligated to the samples with T4 DNA ligase. Samples were washed again with XP Beads, then 100 fmol from them was loaded on a MinION Flow Cell.

#### Direct cDNA-Seq on MinION Flow cells

The same polyA(+) RNA mixture was used for the preparation of the dcDNA library as for dRNA sequencing. The library was generated with the ONT’s direct cDNA sequencing kit (SQK-DCS109) according to the ONT’s protocol. Briefly, the RT reaction was conducted in the presence of anchored oligo(dT) primers (VN primer from the ONT Kit), dNTPs, buffer and enzyme from the Maxima H Minus Reverse Transcriptase kit (Thermo Fisher Scientific), as well as strand switching primer (SSP, including in the ONT kit). To avoid RNA degradation, RNaseOUT™ (Thermo Fisher Scientific) was also added to the reaction. After cDNA synthesis, the RNA strands were digested using RNase Cocktail Enzyme Mix (Thermo Fisher Scientific). The sample was cleaned applying the XP Bead purification method. The second cDNA strand was generated using LongAmp Taq Master Mix from New England Biolabs and the PNT’s PR2 primer. Double-stranded cDNAs were washed with XP Beads then were handled with NEBNext Ultra II End-prep enzyme mix (New England Biolabs). After an XP Bead-washing, sample as ligated to the ONT Adapter Mix using Blunt/TA Ligation Master Mix (New England Biolabs). Finally, 200 fmpl from the XP Bead-purified sample was run on two SpotON Flow Cells.

#### Amplified cDNA-sequencing using MinION device

Total RNA samples from each time point (4, 8, 12, and 20 h p.i.) were sequenced individually using the ONT MinION device and the cDNA-PCR Barcoding protocol (SQK-PCS109 and SQK-PBK004). The RT was carried out as was described at the dcDNA protocol. The samples were amplified using LongAmp Taq Master Mix. Low Input barcode primers (ONT’s SQK-PBK004 kit) were added to the samples with the aim of multiplexing of the samples on the Flow Cells. After the PCR reaction, cDNAs were treated with exonuclease and they were finally cleaned with AMPure XP Beads.

#### Amplified cDNA sequencing using Illumina MiSeq sequencer

The viral transcriptome was also sequenced by using the Illumina SRS approach. The NEXTflex^®^ Rapid Directional qRNA-Seq Kit (PerkinElmer) was used for library preparation from the rRNA-depleted sample. All reagents and enzymes were supplied by the kit and the kit’s manual was followed for the library preparation. RNAs were fragmented enzymatically, then the first and second cDNA strands were generated. Sample was cleaned using AMPure XP Beads, then it was adenylated using the Adenylation Mix from the Kit. Molecular Index Adapters were ligated to the cDNA, then it was washed with XP Beads. Finally, the sample was amplified with PCR and washed with XP Beads. The Illumina MiSeq Reagent Kit v3 (150-cycle format) was used for the sequencing, 12 pM from the library was loaded onto a flow cell.

### Determination of the quantity and quality of RNA samples and sequencing ready libraries

The amount of the purified RNA samples were measured using a Qubit 4 Fluorometer (Thermo Fisher Scientific) and Qubit RNA BR (Broad-Range, quantitation of total RNAs) or Qubit RNA HS Assay Kit (polyA(+) and rRNA-depleted RNAs). The Agilent TapeStation 4150 was used to the determine of the quality of the RNA samples. The amount of the sequencing ready libraries and the quality assessment of the Illumina library was evaluated using the Qubit 4 with Qubit dsDNA High-Sensitivity (HS) and the TapeStation, respectively.

### Data processing and analysis

#### ONT sequencing

Base calling of the raw data from MinION sequencing was carried out with the Guppy software v3.4.5 (ONT). The minimap2 software [48] was used with the following options: -ax splice -Y -C5 –cs to mapping the raw reads to the ASFV genome (accession number: MN715134.1). The LoRTIA (https://github.com/zsolt-balazs/LoRTIA) was applied for the detection of the TSS and TES positions as well as for the determination of full-length reads and transcripts.

The LoRTIA suit was set as follows:

- For dRNA-seq and dcDNA-seq reads: −*5 TGCCATTAGGCCGGG --five score 16 -- check_in_soft 15 3 AAAAAAAAAAAAAAA --three score 16 -spoisson –f True*.
- For o(dT)-primed cDNA reads: −*5 GCTGATATTGCTGGG -- five score 16 --check_in_soft 15 −3 AAAAAAAAAAAAAAA --three_score 16 -spoisson -f True*.

Bam files were visualised using Geneious and IGV tools. The TSSs obtained by Cage-Seq and TESs obtained by p(A)-Seq [29] was mapped to ASFV_HU_2018 genome using Geneious software.

#### Illumina sequencing

The raw Illumina reads were trimmed with the Cutadapt tool [49]. The above-mentioned ASFV reference genome was indexed using STAR aligner v2.7.3a [50] using the following settings: -- genomeSAindexNbases 8, followed by the mapping of the reads with default options. The RPKM and TPM values of individual ORFs were determined using Geneious software (RPKM: reads per kilobase pair per million mapped reads; TPM: transcript per kilobase million).

**Fig. 3.**
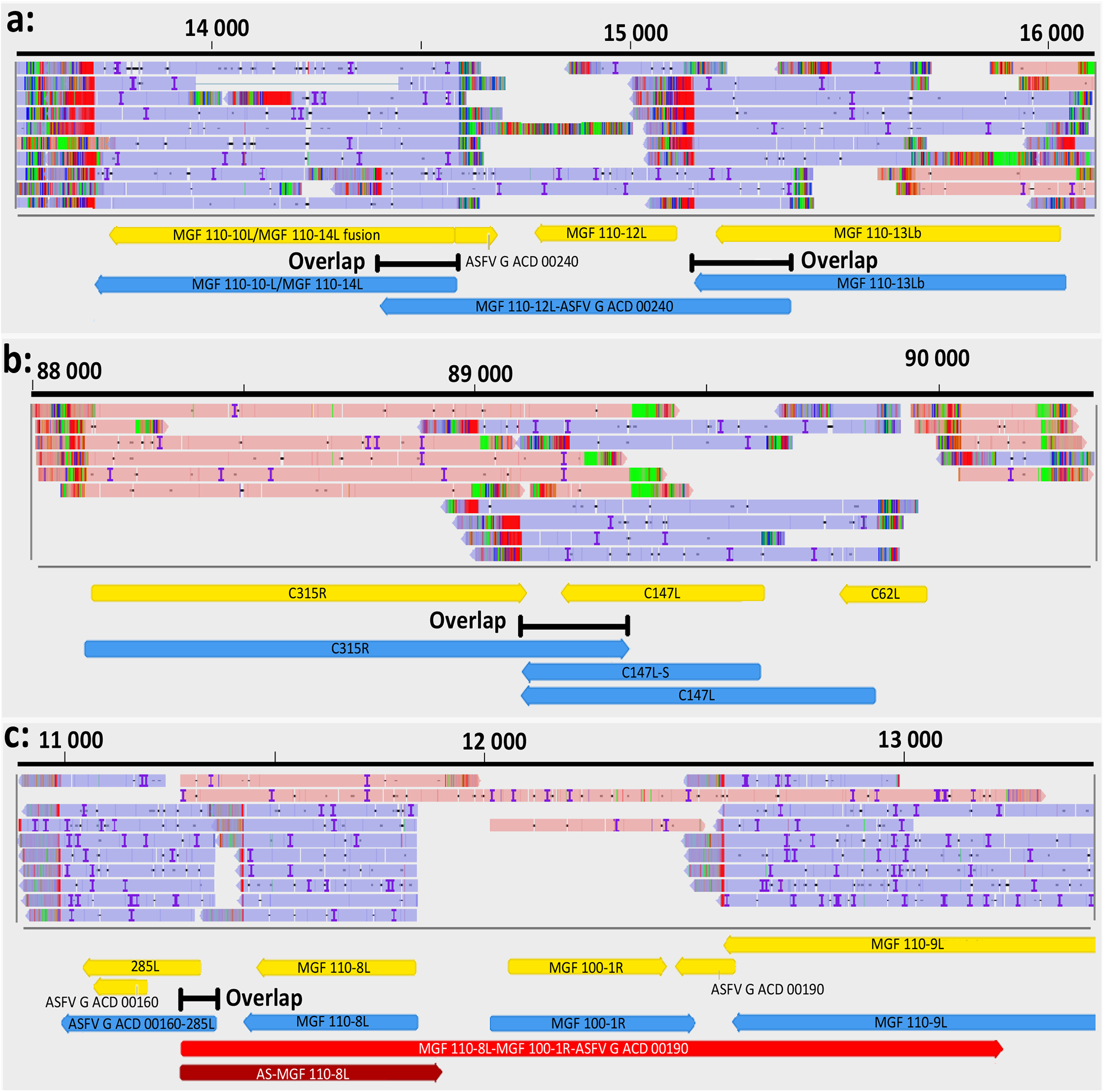
Examples for transcription overlaps of ASFV RNA molecules. (a) parallel overlaps, (b) convergent overlap, (c) Divergent overlap.

## Acknowledgements

This study was supported by the National Research, Development, and Innovation Office grants K 128247 to Z.B. and FK 128252 to D.T. APC was covered by the University of Szeged Open Access Fund: 4636. The funders had no role in study design, data collection and analysis, decision to publish, or preparation of the manuscript.

## Author contributions statement

G.T. generated the ONT dRNA sequencing libraries, and conducted data handling and processing. D.T. performed the MinION sequencing and the Illumina sequencing, participated in data analysis and wrote the manuscript. N.M. carried out bioinformatics analysis. Z.C. conducted MinION direct cDNA sequencing and the Illumina sequencing. I.M. conducted the infection experiments, prepared the pulmonary macrophages and purified the viral total RNA samples. Z.Z. designed the research plan and wrote the manuscript. Z.B. conceived and designed the experiments, managed the study, and wrote the manuscript. All authors read and approved the final paper.

## Additional information

### Accession codes

The LoRTIA software suite is available on GitHub: https://github.com/zsolt-balazs/LoRTIA.

The sequencing datasets generated during this study are available at the European Nucleotide Archive’s SRA database under the accession PRJEB36723.

### Ethics declaration

All methods were performed in accordance with the relevant guidelines and regulations following a protocol approved by the ATK ÁOTI Institutional Animal Care and Use Committee.

### Approvals for animal experiments

Animal were euthanized in the animal facilities of ATK ÁOTI where all methods were performed in accordance with the relevant guidelines and regulations following the protocol approved by the Government Agency of Pest County (PE/EA/1474-7/2017). Approval for the study of ASFV was obtained from the Ministry of Agriculture (Hungary), based on the 189 § (1) Chapter 3, of the 41/1997 (V. 28) Ministerial Decree, about the Veterinary Regulations (Approval number: ÉFHÁT/153-1/2019).

## References

1. Mazur-Panasiuk, N., Żmudzki, J. & Woźniakowski, G. African Swine Fever Virus -Persistence in Different Environmental Conditions and the Possibility of its Indirect Transmission. J. Vet. Res. 2019 63, 303–310; 10.2478/jvetres-2019-0058 (2019)

2. Reis, A. L., Netherton, C. & Dixon, L. K. Unraveling the Armor of a Killer: Evasion of Host Defenses by African Swine Fever Virus. J. Virol. 91, e02338–16; 10.1128/JVI.02338-16 (2017).

3. Achenbach, J. E. et al. Identification of a New Genotype of African Swine Fever Virus in Domestic Pigs from Ethiopia. Transbound. Emerg. Dis. 64, 1393–1404; 10.1111/tbed.12511. (2017).

4. Lubisi, B. A., Bastos, A. D., Dwarka, R. M. & Vosloo, W. Molecular epidemiology of African swine fever in East Africa. Arch. Virol. 150, 2439–52; 10.1007/s00705-005-0602-1. (2005).

5. Olasz, F. et al. The epidemiological features of African swine fever and the possibilities of prevention. Hung. Vet. J., 141, 101–115. (2019).

6. Galindo, I. & Alonso, C. African Swine Fever Virus: A Review. Viruses. 9, 103; doi: 10.3390/v9050103. (2017).

7. Zhou, X. et al. Emergence of African Swine Fever in China, 2018. Transbound. Emerg. Dis. 65, 1482–1484; 10.1111/tbed.12989. (2018).

8. Lu, G. & Zhang, G. African swine fever virus in Asia: Its rapid spread and potential threat to unaffected countries. J. Infect. 80, 350–371; 10.1016/j.jinf.2019.11.011 (2019).

9. Simões, M., Martins, C. & Ferreira, F. Early intranuclear replication of African swine fever virus genome modifies the landscape of the host cell nucleus. Virus Res. 210, 1–7; 10.1016/j.virusres.2015.07.006. (2015).

10. Dixon, L. K., Chapman, D. A. & Netherton, C. L. Upton C. African swine fever virus replication and genomics. Virus Res. 173, 3–14; 10.1016/j.virusres.2012.10.020. (2013).

11. Rodríguez, J. M. & Salas, M. L. African swine fever virus transcription. Virus Res. 173, 15–28; 10.1016/j.virusres.2012.09.014. (2013).

12. Alejo, A., Matamoros, T., Guerra, M. & Andrés, G. A Proteomic Atlas of the African Swine Fever Virus Particle. J. Virol. 92, e01293–18; 10.1128/JVI.01293-18. (2018).

13. Yáñez, R. J. et al. Analysis of the complete nucleotide sequence of African swine fever virus. Virology. 208, 249–78; 10.1006/viro.1995.1149. (1995).

14. Chapman, D. A., Tcherepanov, V., Upton, C. & Dixon, L. K. Comparison of the genome sequences of non-pathogenic and pathogenic African swine fever virus isolates. J. Gen. Virol. 89, 397–408; 10.1099/vir.0.83343-0. (2008).

15. Broyles, S. S. Vaccinia virus transcription. J. Gen. Virol. 84, 2293–303; 10.1099/vir.0.18942-0. (2003).

16. Almazán, F., Rodríguez, J. M., Angulo, A., Viñuela, E. & Rodriguez, J. F. Transcriptional mapping of a late gene coding for the p12 attachment protein of African swine fever virus. J. Virol. 67,553–6; (1993).

17. Almazán, F. et al. Transcriptional analysis of multigene family 110 of African swine fever virus. J. Virol. 66, 6655–67; (1992).

18. Yang Z., Martens, C. A., Bruno, D. P., Porcella, S. F & Moss, B. Pervasive initiation and 3’-end formation of poxvirus postreplicative RNAs. J. Biol. Chem. 287, 31050–60; 10.1074/jbc.M112.390054. (2012).

19. Tombácz, D. et al: Long-Read Isoform Sequencing Reveals a Hidden Complexity of the Transcriptional Landscape of Herpes Simplex Virus Type 1. Front. Microbiol. 8:1079; 10.3389/fmicb.2017.01079. (2017).

20. Balázs, Z. et al. Long-Read Sequencing of Human Cytomegalovirus Transcriptome Reveals RNA Isoforms Carrying Distinct Coding Potentials. Sci. Rep. 7, 15989; 10.1038/s41598-017-16262-z. (2017).

21. Moldován, N. et al. Third-generation Sequencing Reveals Extensive Polycistronism and Transcriptional Overlapping in a Baculovirus. Sci. Rep. 8, 8604; 10.1038/s41598-018-26955-8. (2018).

22. Boldogkői, Z., Moldován, N., Balázs Z., Snyder M. & Tombácz D. Long-Read Sequencing - A Powerful Tool in Viral Transcriptome Research. Trends Microbiol. 27, 578–592 (2019).

23. Tombácz, D. et al. Multiple Long-Read Sequencing Survey of Herpes Simplex Virus Dynamic Transcriptome. Front. Genet. 10, 834; 10.3389/fgene.2019.00834. (2019).

24. Portugal, R. S., Bauer, A. & Keil, G. M. Selection of differently temporally regulated African swine fever virus promoters with variable expression activities and theirapplication for transient and recombinant virus mediated gene expression. Virology. 508, 70–80; 10.1016/j.virol.2017.05.007. (2017).

25. García-Escudero, R, Viñuela, E. Structure of African swine fever virus late promoters: requirement of a TATA sequence at the initiation region. J. Virol. 74, 8176–82; 10.1128/jvi.74.17.8176-8182.2000. (2000).

26. Salas, M. L., Rey-Campos, J., Almendral, J. M., Talavera, A. & Viñuela, E. Transcription and translation maps of African swine fever virus. Virology. 152, 228–40; 10.1016/0042-6822(86)90387-9. (1986).

27. Salas, M. L., Kuznar, J. & Viñuela, E. Polyadenylation, methylation, and capping of the RNA synthesized in vitro by African swine fever virus. Virology. 113, 484–91; 10.1016/0042-6822(81)90176-8. (1981).

28. Jaing, C. et al. Gene expression analysis of whole blood RNA from pigs infected with low and high pathogenic African swine fever viruses. Sci. Rep. 7, 10115; 10.1038/s41598-017-10186-4. (2017).

29. Cackett, G. et al. The African Swine Fever Virus Transcriptome. J. Virol. 94, e00119–20; 10.1128/JVI.00119-20. (2020).

30. Sánchez, E. G. et al. Phenotyping and susceptibility of established porcine cells lines to African Swine Fever Virus infection and viral production. Sci. Rep. 7, 10369; 10.1038/s41598-017-09948-x. (2017).

31. Olasz, F. et al. A Simple Method for Sample Preparation to Facilitate Efficient Whole-Genome Sequencing of African Swine Fever Virus. Viruses. 11, 1129; 10.3390/v11121129. (2019).

32. Sessegolo, C. et al. Transcriptome profiling of mouse samples using nanopore sequencing of cDNA and RNA molecules. Sci. Rep. 9, 14908; 10.1038/s41598-019-51470-9. (2019).

33. Wang, Z., Jia, L., Li, J., Liu, H. & Liu, D. Pan-Genomic Analysis of African Swine Fever Virus. Virol. Sin. 11. 10.1007/s12250-019-00173-6. (2019).

34. Tombácz, D., Balázs, Z., Csabai, Z., Snyder, M. & Boldogkői, Z. Long-Read Sequencing Revealed an Extensive Transcript Complexity in Herpesviruses. Front. Genet. 9, 259; 10.3389/fgene.2018.00259. (2018).

35. Lee, S. et al. Global mapping of translation initiation sites in mammalian cells at single-nucleotide resolution. Proc. Natl. Acad. Sci. U. S. A. 109, E2424–32; 10.1073/pnas.1207846109. (2012).

36. Boldogkői, Z., Balázs, Z., Moldován, N., Prazsák, I. & Tombácz, D. Novel classes of replication-associated transcripts discovered in viruses. RNA Biol. 16, 166–175; 10.1080/15476286.2018.1564468. (2019).

37. Prazsák, I. et al. Long-read sequencing uncovers a complex transcriptome topology in varicella zoster virus. BMC Genomics. 19, 873; 10.1186/s12864-018-5267-8. (2018).

38. Tombácz, D. et al. Dynamic transcriptome profiling dataset of vaccinia virus obtained from long-read sequencing techniques. Gigasci. 7, giy139; 10.1093/gigascience/giy139 (2018).

39. Boldogköi Z. Transcriptional interference networks coordinate the expression of functionally related genes clustered in the same genomic loci. Front. Genet. 3, 122; 10.3389/fgene.2012.00122. (2012).

40. Geballe, A. P. & Mocarski, E. S. Translational control of cytomegalovirus gene expression is mediated by upstream AUG codons. J. Virol. 62, 3334–40 (1988).

41. Calvo, S. E., Pagliarini, D. J. & Mootha, V. K. Upstream open reading frames cause widespread reduction of protein expression and are polymorphic among humans. Proc. Natl. Acad. Sci. 106, 7507–7512; 10.1073/pnas.0810916106 (2009).

42. Barbosa, C. & Romao, L. Translation of the human erythropoietin transcript is regulated by an upstream open reading frame in response to hypoxia. RNA 20, 594–608; 10.1261/rna.040915.113. (2014).

43. Kronstad, L. M., Brulois, K. F., Jung, J. U. & Glaunsinger, B. Dual short upstream open reading frames control translation of a herpesviral polycistronic mRNA. PLoS Pathog. 9, e1003156; 10.1371/journal.ppat.1003156. (2013).

44. Vilela, C. & McCarthy, J. E. G. Regulation of fungal gene expression via short open reading frames in the mRNA 5’untranslated region. Mol. Microbiol. 49, 859–67; 10.1046/j.1365-2958.2003.03622.x (2003).

45. Manual of diagnostic tests and vaccines for terrestrial animals Office International des Epizooties (Paris). Chapitre 3.8.1. Available online: http://www.oie.int/international-standard-setting/terrestrial-manual/access-online/.

46. Tombácz, D. et al. Transcriptome-wide survey of pseudorabies virus using next- and third-generation sequencing platforms. Sci. Data. 5, 180119; 10.1038/sdata.2018.119. (2018).

47. Boldogkői, Z. et al. Transcriptomic study of Herpes simplex virus type-1 using full-length sequencing techniques. Sci. Data. 5, 180266. 10.1038/sdata.2018.266. (2018).

48. Li, H. Minimap2: pairwise alignment for nucleotide sequences. Bioinformatics. 34, 3094–3100; 10.1093/bioinformatics/bty191. (2018).

49. Martin M. Cutadapt removes adapter sequences from high-throughput sequencing reads. EMBnet.Journal. 17, 10–12; 10.14806/ej.17.1.200. (2011).

50. Dobin, A. et al. STAR: ultrafast universal RNA-seq aligner. Bioinformatics. 29, 15–21; 10.1093/bioinformatics/bts635 (2013).

